# Outwitting planarian’s antibacterial defence mechanisms: *Rickettsiales* bacterial trans-infection from *Paramecium multimicronucleatum* to planarians

**DOI:** 10.1101/688770

**Authors:** Letizia Modeo, Alessandra Salvetti, Leonardo Rossi, Michele Castelli, Franziska Szokoli, Sascha Krenek, Elena Sabaneyeva, Graziano Di Giuseppe, Sergei I. Fokin, Franco Verni, Giulio Petroni

## Abstract

Most of the microorganisms belonging to genera responsible for vector-borne diseases (VBD) have hematophagous arthropods as vector/reservoir. Recently, many new species of microorganisms phylogenetically related to agents of VBD were found in a variety of aquatic eukaryotic hosts, in particular, numerous new bacterial species related to the genus *Rickettsia* (*Alphaproteobacteria, Rickettsiales*) were discovered in protist ciliates and other unicellular eukaryotes. Although their pathogenicity for humans and terrestrial animals is not known, these bacteria might act as etiological agents of possible VBD of aquatic organisms, with protist as vectors. In the present study, we characterized a novel strain of the *Rickettsia*-Like Organism (RLO) endosymbiont “*Candidatus* (*Ca*.) Trichorickettsia mobilis” in the macronucleus of the ciliate *Paramecium multimicronucleatum* through Fluorescence *In Situ* Hybridization (FISH) and molecular analyses. Ultrastructural investigations on the presence of flagella confirmed previous studies on the same bacterial species. The potential trans-infection per *os* of this bacterium to planarians (*Dugesia japonica*), a widely used model system able to eliminate a wide range of bacteria pathogenic to humans and other Metazoa, was further verified. Ciliate mass cultures were set up, and trans-infection experiments were performed by adding homogenized paramecia to food of antibiotic-treated planarians, performed. Treated and non-treated (i.e. control) planarians were investigated at day 1, 3, and 7 after feeding for endosymbiont presence by means of PCR and ultrastructural analyses. Obtained results were fully concordant and suggest that this RLO endosymbiont can be transferred from ciliates to metazoans, being detected up to day 7 in treated planarian enterocytes inside and, possibly, outside phagosomes.

## Introduction

Bacteria of order *Rickettsiales* (*Alphaproteobacteria*) live in an obligate association with a wide range of eukaryotes [1–8], and, for their vast majority, are localized intracellularly, although the case of an extracellular *Rickettsiales* bacterium was recently documented [9]. *Rickettsiales* are widely studied for their involvement in medical and veterinary fields. Indeed, many of them (e.g. *Rickettsia* spp., *Anaplasma* spp., *Ehrlichia* spp., and *Orientia tsutsugamushi*) are vectored by lice, ticks and mites, and cause mild to severe disease [1, 10], such as epidemic typhus, Rocky Mountain spotted fever [11–13], anaplasmosis, ehrlichiosis [14, 15], heartwater [16], and scrub typhus [17].

In the last two decades, many new genera and species of *Rickettsiales* were found as symbionts in a variety of other non-vector eukaryotic hosts, both from terrestrial and aquatic environments [reviewed in 3, 4, 7]. In particular, numerous such novel bacterial species were retrieved in aquatic protists [18–22], including, notably, parasitic (*Ichthyophthirius multifiliis*) [23, 24] and various free-living ciliates (e.g. *Paramecium* spp., *Euplotes* spp., *Diophrys oligothrix, Pseudomicrothorax dubius, Spirostomum minus*, and *Colpidium striatum*) [25–33].

Such findings have clearly indicated the ability of those “aquatic” *Rickettsiales* to perform horizontal transmission including host species shift, although mostly from indirect evidence, i.e. closely related bacteria in hosts as different as ciliates, hydra [34], corals [35], ring worms [36], ascidians [37, 38], as well as incongruent host and symbionts phylogenies [3, 39, 40]. In few cases, direct evidence in experimental interspecific transfers between unicellular hosts was produced [19, 29].

In most cases the relationships between *Rickettsiales* associated with aquatic eukaryotes and their hosts were not clarified in detail, as in general no evident effect on host biology was observed [7], with the exception of “*Candidatus* (*Ca*.) Xenohaliotis californiensis”, which is considered the cause of the withering syndrome in its abalone hosts [41, 42]. It has been hypothesized by different authors that bacteria, possibly including *Rickettsiales*, harboured by aquatic protists might constitute etiological agents of possible Vector Borne Diseases (VBD) of aquatic animals [55, 61]. Indeed, protists could be the potential vectors for some of the numerous cases of epidemics caused by *Rickettsia*-Like Organisms (RLOs; i.e. intracellular bacteria morphologically similar to *Rickettsia*) determining an increasing number of massive deaths in intensive aquaculture facilities during the last years [62], including molluscs [41, 43, 44], crustaceans [45–47] and fishes [48–50]. Although many RLOs were actually shown to be phylogenetically unrelated to *Rickettsiales* (e.g. *Gammaproteobacteria*) [51, 52, 46], at least in one case available data indicate a probable connection between a truly *Rickettsiales* bacterium and fish disease. In detail, a “*Ca*. Midichloria mitochondrii”-related bacterium was linked to the red-mark syndrome in rainbow trout [53–54]. Most importantly, although a transmission route was not directly proven, the same bacterium was found in association with the ciliate *I. multifiliis* [24], which is indeed a fish parasite. Interestingly, even other *Rickettsiales* symbionts (“*Ca*. Megaira” genus) can be found in the same ciliate host species, which might suggest a potential transmission route for these symbionts as well [23, 24].

Taking into account such premises, in order to investigate the hypothesis that protists can act as natural reservoir for potentially pathogenic bacteria [25, 55–61], we experimentally tested for the first time the possibility that endosymbionts of ciliates could be transferred to aquatic Metazoa. The *Rickettsiales* “*Ca*. Trichorickettsia” was chosen as candidate for trans-infection experiments as it shows a broad ciliate host range, infecting different cell compartment (i.e. cytoplasm and nucleus) of *Paramecium multimicronucleatum, Paramecium nephridiatum, Paramecium calkinsi* and *Euplotes aediculatus* [28, 63]. Additionally, in *P. multimicronucleatum* and *P. calkinsi*, “*Ca*. Trichorickettsia” appears covered by long flagella and is able to actively move [28, 63]. Thus, for our research purpose, we selected the newly isolated *P. multimicronucleatum* strain US_Bl 16I1 as donor host in the trans-infection experiments. We characterized its macronuclear bacterial symbiont, assigning it to “*Ca*. Trichorickettsia mobilis subspecies hyperinfectiva” (*Rickettsiaceae, Rickettsiales*), previously discovered and described in the cytoplasm of *Paramecium calkinsi* [63]. As recipient, we selected the freshwater planarian *Dugesia japonica*. Freshwater planarians are benthic organisms, living in mud and under rocks in ponds and streams. They are zoophages or fleshing-eating animals, but ingest also detritus, fungi, and bacteria [64]. Thus, they may encounter in their environment a large variety of microbes [65], including ciliates possibly hosting endosymbionts. Planarians have always been considered an important model for studying stem cells and regeneration [66], but recently they became also important for studying the natural immunity system of Metazoa [67–72]. Based on all these considerations, they appeared a suitable model for such experiments.

Bacterial endosymbiont trans-infection from *P. multimicronucleatum* to planarians was investigated by checking for the presence of “*Ca*. Trichorickettsia mobilis” in tissues of ciliate-treated planarians by means of PCR and Transmission Electron Microscopy (TEM). The findings obtained by means of the two investigation approaches allowed us to test the hypothesis that this endosymbiont can transfer from ciliate protists to Metazoa.

## Materials and methods

### Ciliate host isolation, culturing, and identification

*Paramecium multimicronucleatum* monoclonal strain US_Bl 16I1 was established from a cell isolated from a freshwater sample collected from the Yellowwood Lake (39°11’29,0’’N, 86°20’31,4’’W), IN, USA and cultivated in San Benedetto mineral water (San Benedetto S. p. A., Italy)

The strain was maintained in the laboratory in an incubator at a temperature of 19 ± 1 °C and on a 12:12 light/dark cycle (irradiance by means of NATURAL L36W/76 and FLORA L36W/77 neon tubes, OSRAM, Berlin, Germany). Instead of using bacteria as common food source for paramecia [73], cells were fed on monoclonal cultures of flagellated green algae, i.e. *Chlorogonium* sp. (freshwater) or, alternatively, *Dunaliella tertiolecta* (brackish water, 1 ‰ of salinity) to minimize bacterial load in cell cultures. They were fed two to three times per week to obtain a mass culture (1.5 L) suitable for trans-infection experiments. Living ciliates were observed with a Leitz (Weitzlar, Germany) microscope equipped with differential interference contrast (DIC) at a magnification of 100-1250x for general morphology and swimming behaviour according to [74, 75]. DIC observation of ciliates was also aimed at checking for macronuclear endosymbiont motility. Feulgen stained ciliates were observed to obtain information on the nuclear apparatus. Microscope images were captured with a digital camera (Canon PowerShot S45). Cell measurements were performed using ImageJ (https://imagej.nih.gov/ij/). Morphological identification was conducted according to literature data [76].

### Characterization of ciliate and endosymbiont

Characterization of the ciliate host and its bacterial endosymbiont was performed through the “full cycle rRNA approach” [77], i.e. through the nuclear 18S rRNA gene (generally considered the preferred marker to study molecular taxonomy and phylogeny of eukaryotic organisms - just as an example see [78]), ITS region and partial 28S gene (ciliate), and 16S rRNA gene (endosymbiont) sequencing, combined with Fluorescence *In Situ* Hybridization (FISH) experiments. Additionally, mitochondrial cytochrome c oxidase subunit I (COI) gene was sequenced as well, to display haplogroup variation in *P. multimicronucleatum* [79].

#### Molecular characterization

For molecular characterization of both ciliate host and bacterial endosymbiont, total DNA was extracted from approximately 50 *Paramecium* cells as described in [30, 31]. All PCR reactions were carried out with the Takara ExTaq (Takara, Kusatsu, Japan) reaction kit. COI gene was amplified using the degenerated forward primer F338dT and reverse primer R1184dT [80], while for 18S rRNA gene amplification the forward 18S F9 Euk [81] and the reverse 18S R1513 Hypo primers [82] were used. The ITS region (including partial 18S rRNA gene, ITS1, 5.8S rRNA gene, ITS2, and partial 28S rRNA gene) sequence was obtained by PCR with the forward 18S F919 and the reverse 28S R671 primers (modified according to [83]). 16S rRNA gene of the bacterial symbiont was amplified with the *Alphaproteobacteria*-specific forward primer 16S Alpha F19b and the universal bacterial reverse primer 16S R1522a [84]. Purified PCR products by NucleoSpin gel and PCR cleanup kit, Macherey-Nagel GmbH Düren, Germany) were directly sequenced at GATC Biotech AG (Constance, Germany). Internal primers were used to sequence 16S rRNA gene [84], 18S rRNA gene [85] and ITS regions [83]. The latter two sequences were then joined together, exploiting the partial overlap on the 18S rRNA gene sequences. For the COI gene sequencing, amplification primers were employed from both directions.

#### FISH experiments

Preliminary FISH experiments were performed using the eubacterial universal probe EUB338 [86] labelled with fluorescein-isothiocyanate (FITC) and the specifically designed probe Rick_697 (5’-TGTTCCTCCTAATATCTAAGAA-3’) labelled with Cy3 to verify the presence of endosymbiotic bacteria belonging to the family *Rickettsiaceae* [28]. Based on the obtained results, i.e. the presence of a single positive signal in the ciliate macronucleus and the newly characterized 16S rRNA gene sequence corresponding to “*Ca*. Trichorickettsia mobilis subspecies hyperinfectiva”, a second FISH experiment was carried out using a species-specific probe, i.e. the probe Trichorick_142 (5’-GTTTCCAAATGTTATTCCATAC-3’) [28] in combination with the eubacterial universal probe EUB338 [86]. The experiments followed the protocol by [87] employing a hybridization buffer containing no formamide, according to the recommendations for the used probes. Slides were mounted with SlowFade Gold Antifade with DAPI (Invitrogen, Carlsbad, USA) and viewed with a Zeiss AxioPlan fluorescence microscope (Carl Zeiss, Oberkochen, Germany) equipped with an HBO 100W/2 mercuric vapor lamp. Digital images were captured by means of a dedicated software (ACT2U, version 1.0).

### Planarian culturing

Planarians used in this work belonged to the species *Dugesia japonica*, asexual strain GI [88]. Animals were kept in artificial water (CaCl_2_ 2.5mM; MgSO_4_ 0.4mM; NaHCO_3_ 0.8mM; KCl 77μM;) at 18°C in dim light conditions and fed with chicken liver (purchased from local food stores) prepared according to [89], once a week. Non-regenerating specimens, within 5-8 mm of length, were used for all experimental procedures, after being starved for about two weeks.

### Transmission electron microscopy (TEM)

TEM preparations were obtained both for *P. multimicronucleatum* cells and planarians to study the ultrastructure of the ciliate-associated endosymbiont “*Ca*. Trichorickettsia mobilis subspecies hyperinfectiva” and to verify the success of trans-infection experiments (i.e. to check the animals for the presence of trans-infected ciliate endosymbionts in their tissues) respectively.

TEM preparations of *P. multimicronucleatum* cells were obtained according to [90]. Briefly, cells were fixed in a 1:1 mixture of 2.5% (v/v) glutaraldehyde in 0.1 M cacodylate buffer (pH 7.4), and 2% (w/v) OsO_4_ in distilled water, ethanol and acetone dehydrated, and embedded in Epon-Araldite mixture. Ultrathin sections were stained with 4% (w/v) uranyl acetate followed by 0.2% (w/v) lead citrate.

TEM preparations of planarian specimens were obtained as previously described [91, 92], with minor modifications. Therefore, planarians were fixed with 3 % glutaraldehyde in 0.1 M cacodylate buffer, and post-fixed with 2 % osmium tetroxide for 2 h. After ethanol dehydration, samples were embedded in Epon-Araldite mixture. Ultrathin sections were stained with uranyl acetate and lead citrate and observed with a Jeol 100 SX transmission electron microscope.

#### Negative staining

For negative staining, several *P. multimicronucleatum* cells of the strain US_Bl 16I1 were washed in distilled water and squashed by means of a syringe; a drop of the resulting suspension was placed on a grid. Bacteria were allowed to precipitate for 2–3 min, then a drop of 1% uranyl acetate in distilled water was added for no longer than 1 min. The liquid was then absorbed with filter paper, the grid was air-dried, and observed under TEM.

### Trans-infection experiments

Two trans-infection experiments with the same protocol were sequentially carried out along a period of four months. Each experiment was conducted by treating a fixed number of planarians with ciliate-enriched food, i.e. liver paste mixed with *P. multimicronucleatum* homogenate; from now on these animals will be referred to as “treated planarians”. As for experimental control, planarians fed on plain liver paste were used; these animals will be referred to as “control planarians”. Each trans-infection experiment was performed according to the following protocol:

1. 96 planarians were selected by eye from culture (see above), washed in fresh culturing water and collected in a Petri dish with 50 μg/ml gentamicin sulfate (Sigma, Saint Luis, MO, USA) dissolved in their culturing water (see above). This preliminary antibiotic treatment was performed to minimize potential endogenous bacteria contaminants without endangering the animals [93]. Planarians were left in antibiotic treatment for 24 h under regular culturing conditions concerning temperature and light (see above) and kept under visual control during that period to verify their viability during the antibiotic treatment and by the time of trans-infection. Then, planarians were washed 6 times in their fresh culturing water and split into two equal groups of 48 individuals each and accommodated in two different Petri dishes for the trans-infection procedure by means of feeding.
2. 1.5 L of *P. multimicronucleatum* mass culture (cell concentration: ~ 4 x 10^5^ cell/L) was filtered with a nylon filter (pore size: 100 μm). Cells were washed twice in sterile San Benedetto mineral water to minimize potential bacterial contaminants, concentrated and harvested by means of centrifugation (400 x *g* per 10 min), so to reduce the medium volume to 2 ml. Then, cells were mechanically homogenized for 20 min by repeated passages through a syringe (needle: 22GA, 0.70 mm in diameter). Cell homogenate was centrifuged (10,000 x *g* per 10 min), and the resulting pellet was resuspended in 50 μl of planarian food (homogenized liver paste) by direct resuspension.
3. A group of 48 planarians was fed on *P. multimicronucleatum-enriched* liver paste (treated planarians), while the other group was in parallel fed on plain liver paste (control planarians). For feeding, food was sown on the bottom of the Petri dish. Animals were allowed to reach it and comfortably feed for a period of 2h under regular culturing conditions (see above). Attention was paid to planarian feeding behaviour during this period. As no differences were noted concerning feeding behaviour between treated and control planarians, and feeding procedure was exerted by all planarians as expected according to regular planarian culturing, we proceeded with the next steps. Two washing steps were then carried out removing the medium and adding fresh planarian culturing water. Finally, the two groups of animals were left in their fresh culturing water and in regular cultivation conditions in the two Petri dishes until the collection of specimens for the next TEM and PCR analyses at the three timepoints (see below) of the experiments. Animals were kept under visual control throughout the experiment to regularly check their viability.
4. At day 1, 3, and 7 after feeding (experimental timepoints), from each of the two Petri dishes containing treated and control planarians a group consisting of 16 animals were sampled: 4 animals were fixed and processed for TEM, and 12 were rapidly frozen and stored at −80°C and dedicated to DNA extraction (4 animals per each sample).

### PCR verification of trans-infection success

Treated and control planarians were processed for genomic DNA extraction, purification, and PCR amplification with endosymbiont-specific primers. For each experimental condition (treated and control planarians) and each experimental timepoint (1, 3, and 7 days after feeding), genomic DNA was extracted from frozen samples by using the Wizard Genomics DNA purification kit (Promega, Madison, WI, USA). One microliter of purified DNA was analysed by gel electrophoresis to check for integrity. DNA was quantified using a Nanodrop spectrophotometer. For each experimental timepoint of both experimental conditions (i.e. treated and control planarians), similar amounts of genomic DNA were used for amplification using the ampli-Taq-gold DNA polymerase (Applied Biosystems, Foster City, CA, USA) and for a nested PCR assay. The first primer pair used, specific for “*Ca*. Trichorickettsia mobilis”, was RickFla_ F69 5’-GTTAACTTAGGGCTTGCTC-3’ and Rick_R1455 5’-CCGTGGTTGGCTGCCT-3’ [28]; PCR conditions were: 3 minutes at 94°C; 5 cycles each consisting of 30 s at 94°C, 30 s at 58°C, 2 minutes at 72°C; 10 cycles each consisting of 30 s at 94°C, 30 s at 54°C, 2 minutes at 72°C; 25 cycles each consisting of 30 s at 94°C, 30 s at 50°C, 2 minutes at 72°C; ending with 7 minutes at 72°C. The second, nested primer pair used was RickFla_F87 5’-CTCTAGGTTAATCAGTAGCAA-3’ and Rick_R1270 5’-TTTTAGGGATTTGCTCCACG-3’ [28]. For nested PCR assay, one microliter of PCR product of the first PCR assay was used as a template; PCR conditions were as above.

For each experimental condition and each timepoint of the two trans-infection experiments, the DNA amplification was performed in duplicate. Samples were considered positive if a single band of the expected size was recorded after nested amplification. Sequencing of amplicons was carried out to confirm the presence of “*Ca*. Trichorickettsia mobilis” using the primer RickFla_F87 (see above) and Sanger sequencing (BMR Genomics, Padova, Italy).

Samples obtained from control planarians were used as PCR negative control. As positive control, genomic DNA purified from *P. multimicronucleatum* monoclonal strain US_Bl 16I1 was processed.

## Results

### Host morphological and molecular identification

Ciliate strain US_Bl 16I1 (Figs 1a, 1b; Supplementary Material; S1 and S2 Figs) was confirmed in morphological inspections as *Paramecium multimicronucleatum* Powers and Mitchell, 1910 [94] considering features of key-characters such as cell size, number and features of micronuclei (mi), and number and features of contractile vacuoles (CV) (S1 and S2 Figs), as described in previous literature [76, 95–98]. The molecular analysis of the combined (partial) 18S rRNA-ITS1-5S rRNA gene-ITS2-(partial) 28S rRNA gene sequence (2792 bp, GenBank accession number: MK595741) confirmed the species assignation by morphological identification with a 100% sequence identity with the sequences of other *P. multimicronucleatum* strains already present in GenBank presenting either 18S rRNA gene portion only (AB252006 and AF255361), or the ITS portion (AY833383, KF287719, JF741240 and JF741241). COI gene sequence identity of strain US_Bl 16I1 (760 bp, accession number: MK806287) is highest with another *P. multimicronucleatum* haplotype (FJ905144.1; 96.3%).

**Fig 1.**
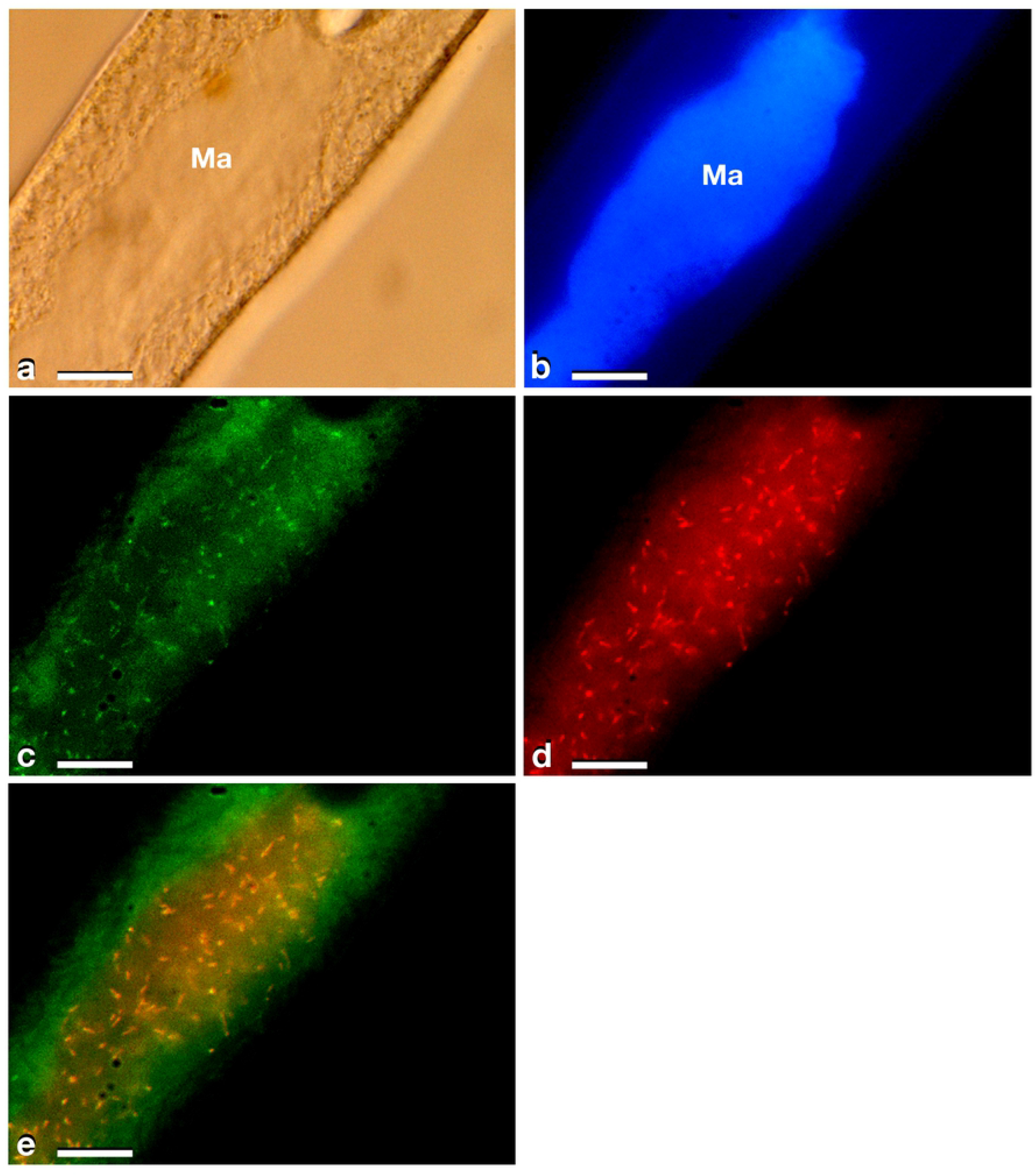
Results of FISH experiments on a *P. multimicronucleatum* US_Bl 16I1 cell. (**a**) bright field microscopy (Ma, macronucleus). (**b**) after DAPI staining. (**c**) treated with the almost universal bacterial probe EUB338 double labelled with fluorescein (green) (probe labelling both at 5’ and 3’ ends). (**d**) treated with species-specific probe Trichorick_142 labelled with Cy3 (red) targeting “*Ca*. Trichorickettsia mobilis”. (**e**) merge of **c** and **d**. The number of endosymbionts targeted by the species-specific probe in the macronucleus reached ~ 100 (**d, e**). Scale bars stand for 10 μm.

### Endosymbiont identification and features

As in *Paramecium* three different subspecies of RLOs “*Ca*. Trichorickettsia mobilis” have been described [63], 16S rRNA gene amplification was performed to identify endosymbionts associated with *P. multimicronuclatum* US_Bl 16I1 strain (sequence length 1,563 bp, accession number: MK598854), which allowed to assign it to “*Ca*. Trichorickettsia mobilis subsp. hyperinfectiva”, an endosymbiont described in the cytoplasm of *Paramecium calkinsi* (99.8 % identity of the novel sequence with type strain MF039744.1) [63]. FISH experiments (Figs 1c-1e) performed by using eubacterial universal probe EUB338 (Fig 1c) and the species-specific probe Trichorick_142 (Fig 1d) confirmed this result but disclosed a different, novel bacterial localization, i.e. the ciliate macronucleus (roughly with a presence of about 100 endosymbionts). Additionally, the full overlapping of eubacterial universal probe EUB338 and “*Ca*. Trichorickettsia”-specific probe signals indicated that this symbiont constituted the total set of intracellular bacteria in host *P. multimicronucleatum* US_Bl 16I1 cells (Fig 1e, merge).

In TEM-processed ciliate cells, the endosymbionts were confirmed to be hosted in the macronucleus; they showed a two-membrane cell wall typical for Gram-negative bacteria and appeared rod-shaped, with rounded to narrower ends (Fig 2). They were surrounded by a clear halo and not encircled by any symbiosomal vacuole, i.e. they appeared in direct contact with host nuclear material. Their size was ~ 1.2--2.1 x 0.5--0.6 μm, and they were scattered throughout the macronucleus of the ciliate (Figs 2a-2d). Sometimes in their cytoplasm several electron-lucid “holes” (diameter: ~ 0.2 μm), were observed (Fig 2a), as already described in the same endosymbiont in *P. calkinsi* [63]. Bacterial cytoplasm was electrondense and rich in ribosomes (Fig 2b); the presence of additional cytoplasmic components (e.g. electron-dense particles with a special arrangement) was not disclosed. Endosymbionts displayed thin (diameter: ~ 0.009 μm) and short flagella distributed all around the cell (Figs 2a and 2c-2d) which sometimes formed a putative tail emerging from a cell end (Fig 2d). The presence of longer flagella was evident after negative staining procedure (Fig 3): besides some short flagella occurring all around the cell, at least a few longer and thicker flagella (~ 2 x 0.012 μm) arose from one of cell ends.

**Fig 2.**
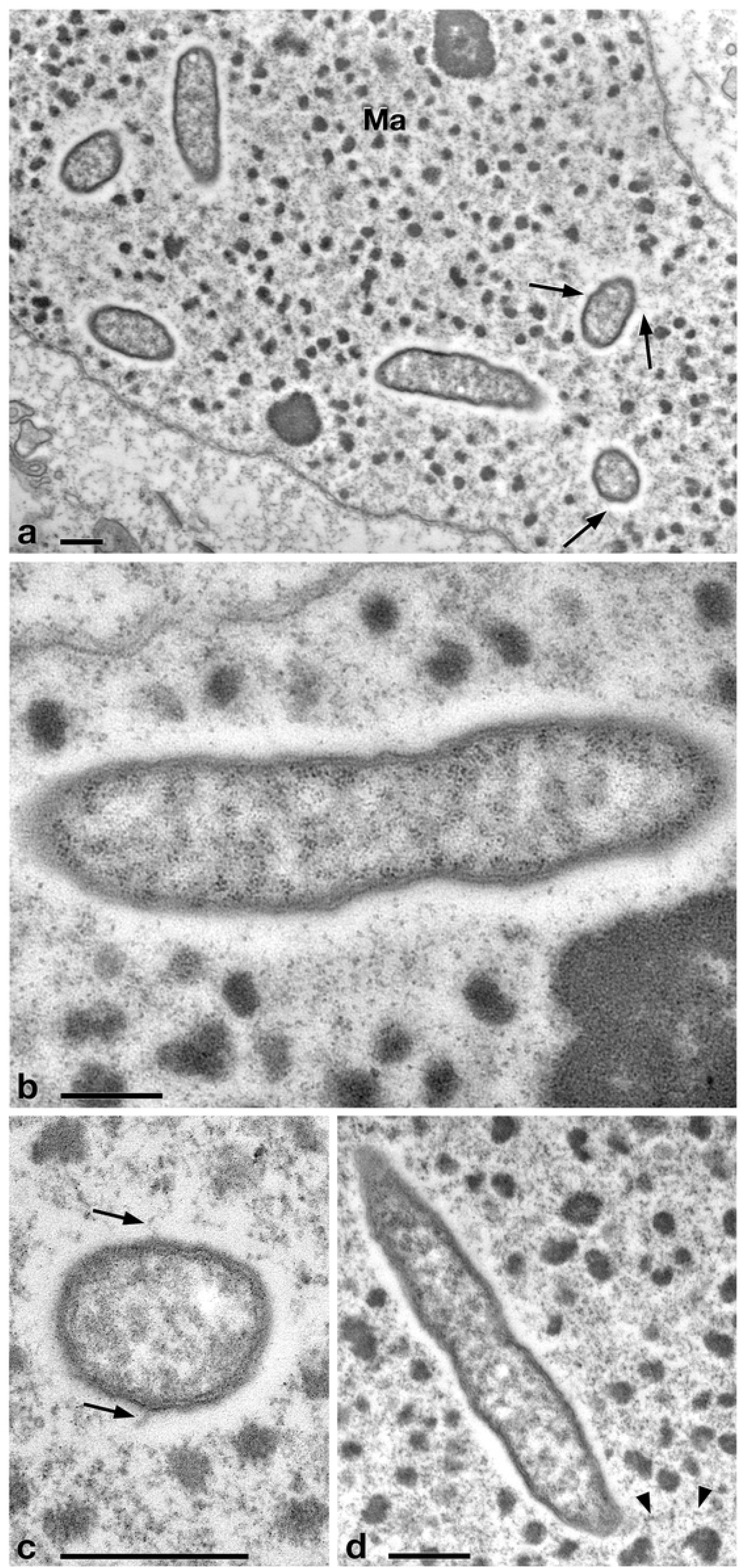
TEM pictures showing the morphological-ultrastructural details of “*Ca*. Trickorickettsia mobilis subsp. hyperinfectiva” in the macronucleus of *P. multimicronucleatum* US_Bl 16I1. (**a**) longitudinally and transversely-sectioned endosymbionts inside macronucleus (Ma) with a double membrane and surrounded by a clear halo; electron-lucid roundish areas occur in the cytoplasm of some endosymbionts; (**b**) numerous bacterial ribosomes are visible; (**c, d**) bacterial flagella are short, located either around (arrow) the endosymbiont cell or at a cell pole, where they can form a putative tail (arrowhead). Scale bars stand for 0.5 μm.

**Fig 3.**
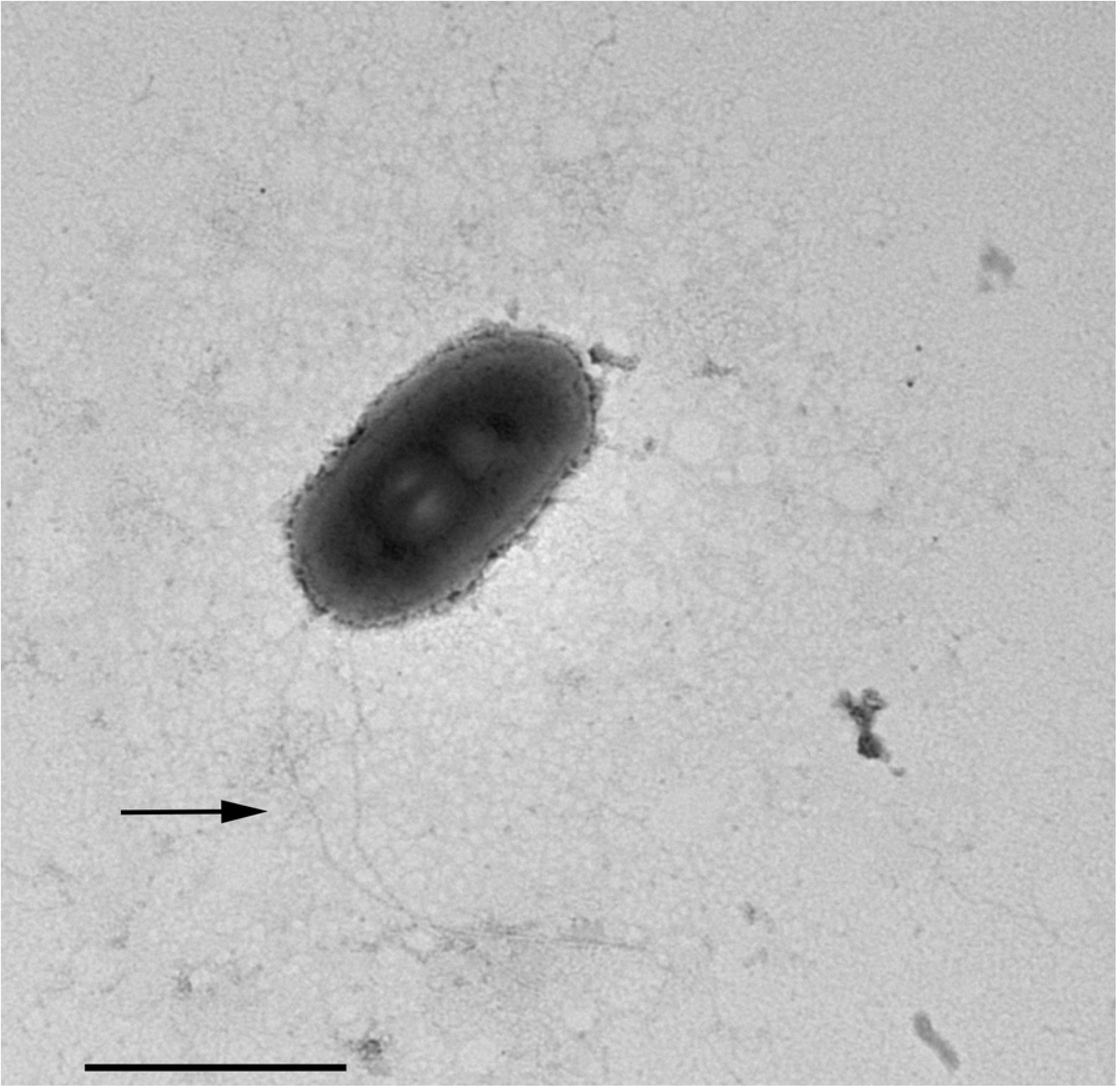
Negative contrast at TEM of a specimen of “*Ca*. Trickorickettsia mobilis subsp. hyperinfectiva”. Some short flagella are present and at least two longer flagella (arrow) arise from one of cell ends. Scale bar stands for 1 μm.

Some active bacterial motility, likely obtained by means of their flagella, was recorded inside the macronucleus of intact *P. multimicronucleatum* cells during observation under DIC microscope at 1,000x. Endosymbionts were seen to move through the chromatin bodies covering short distances (Supplementary Movie SM1). After ciliate cell squashing the released bacteria were still capable of movement (data not shown).

### Trans-infection experiments

The success of each of the two independent trans-infection experiments performed was investigated by means of DNA extraction/PCR experiments and TEM observation on treated (i.e. fed on ciliate homogenate added to liver paste) and control (i.e. untreated, fed on plain liver paste) planarians at the different experimental timepoints, according to the described experimental procedure. By verifying the possible presence of *P. multimicronucleatum* endosymbiont “*Ca*. Trichorickettsia mobilis” in extracted as well as TEM processed animals, the two investigation methods produced completely overlapping results on planarians used in the two trans-infection experiments and with full concordance on trans-infection success. Thus, for each of the two different investigation methods, results of the two trans-infection experiments are concurrently reported in the following sections. Similarly, figures showing the results of the trans-infection experiment at the different experimental timepoints (Figs 4 and 5, S3 Fig) are meant to be representative for both performed experiments, irrespective of the specific experiment they refer to.

**Fig 4.**
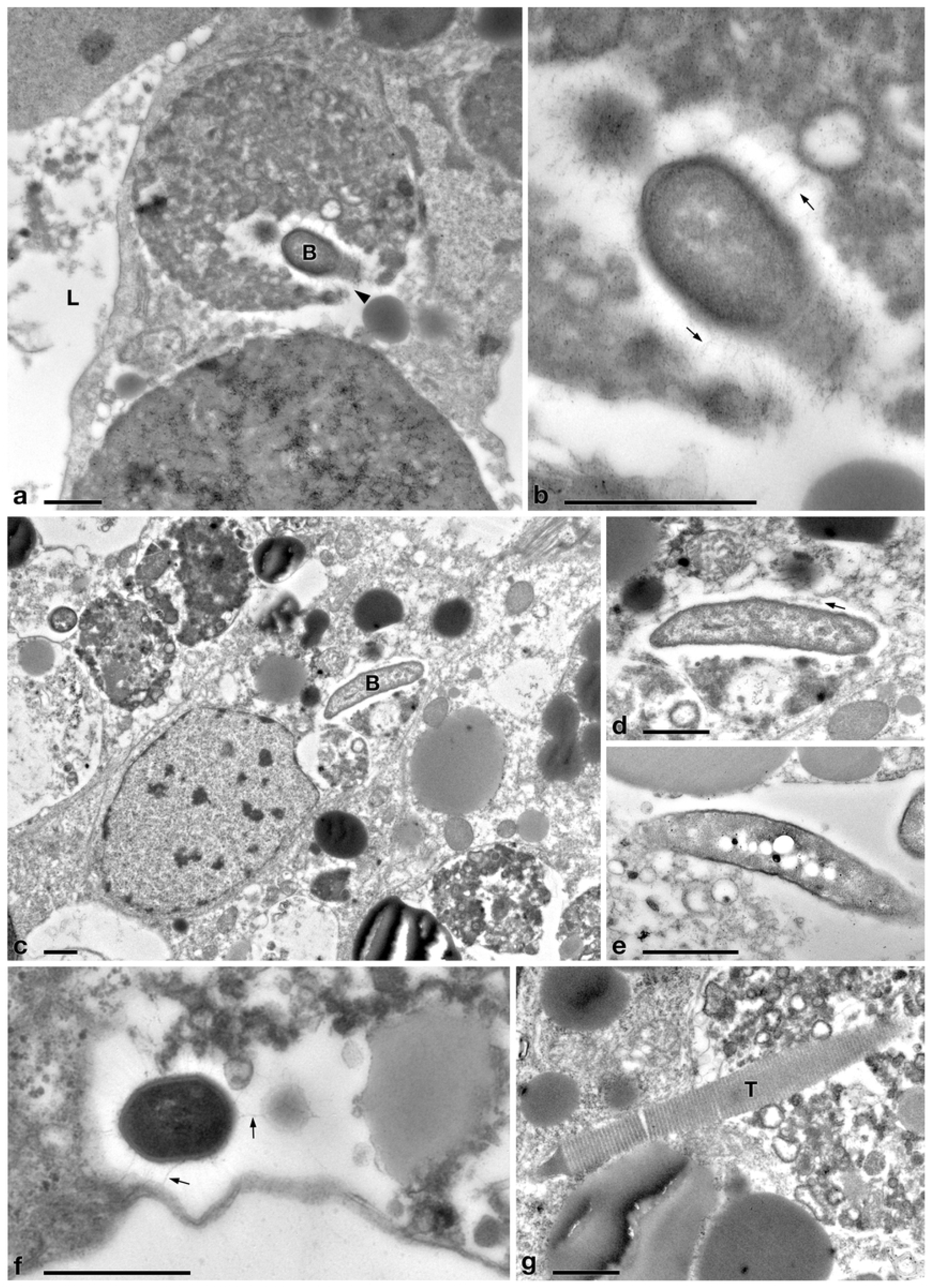
TEM pictures of the intestine of *D. japonica* treated with liver paste enriched with the homogenate of *P. multimicronucleatum* US_Bl 16I1 cells and fixed at day 1 after feeding. (**a, b**) (**b**, enlargement of a particular of **a**) a flagellated bacterium (B) inside a phagosome whose membrane is interrupted (arrowhead); L, lumen of planarian intestine; (**c, d**) (**d**, enlargement of a particular of **c**) a longitudinally sectioned bacterium (B) free in the cytoplasm of an intestinal cell of a treated planaria; (**e**) another free bacterium, showing cytoplasmic electron-lucid “holes”; (**f**) a recovered flagellated bacterium in cross section. (**g**) an intact extruded trichocyst (T) (extrusive organelle) of *Paramecium* detected in planarian intestine. Arrows, flagella. Scale bars stand for 1 μm (**a-e, g**), and 0.5 μm (**f**).

**Fig 5.**
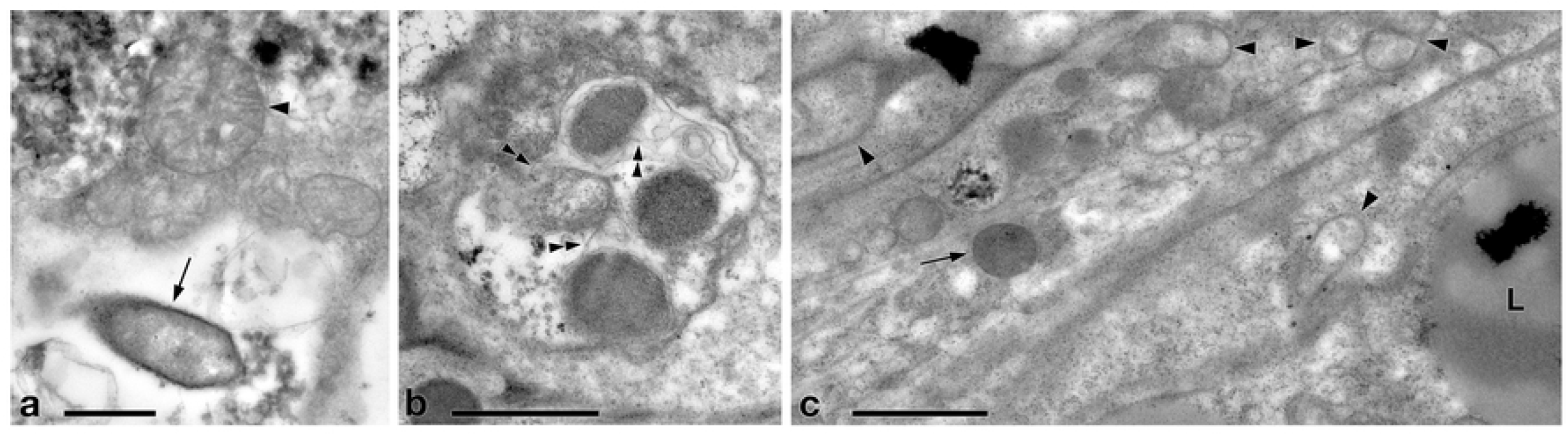
TEM pictures of the intestine of *D. japonica* fixed at day 7 after feeding with liver paste enriched with the homogenate of *P. multimicronucleatum* US_Bl 16I1 cells. (**a, b**) bacteria recognizable inside phagosomes and (**c**) bacterial shapes in the cytoplasm of an intestinal cell. L, lipid droplet; arrow, bacteria; arrowhead, mitochondria; double arrowhead, flagella. Scale bars stand for 1 μm.

#### PCR experiments

The DNA of “*Ca*. Trichorickettsia mobilis” was detected at all experimental timepoints (1, 3, and 7 days after feeding) in genomic DNA preparations obtained from all investigated planarians fed on *P. multimicronucleatum* lysate-enriched liver paste. The size of the nested amplicon (1360 bp) obtained from treated planarian samples matched the size of the amplicon obtained in DNA purified from ciliate monoclonal strain US_Bl 16I1 (positive control). The sequencing of the obtained amplicons confirmed the specificity for “*Ca*. Trichorickettsia mobilis” (100% sequence identity with MF039744.1).

No amplification product was obtained in genomic DNA samples from control planarians fed on plain liver paste (negative control).

#### TEM observation

Ultrastructural observation was conducted on TEM-processed specimens per each timepoints of the experiments, for both experimental groups, i.e. treated and control planarians.

In tissues collected from the investigated planarians fed on *Paramecium* US_Bl 16I1 lysate-enriched liver paste, the presence of bacteria with morphology and sizes (see below for details) fitting with those of ciliate endosymbionts within intestinal cells was detected at all timepoints of the two experiments (Figs 4 and 5).

One day after feeding, several bacteria were recognizable in planarian enterocyte phagosomes. Some of them were still intact (Figs 4a and 4b), even performing cell division. Others appeared degraded, being subjected to digestion (data not shown). In several cases, membrane of bacteria-including phagosomes was damaged and interrupted (Figs 4a and 4b). Bacteria free in the cytoplasm of planarian intestinal cells, i.e. in direct contact with their cytoplasm, were detected as well; they showed a two-membrane cell wall and a surrounding clear halo (Figs 4c-4f), with electron-lucid “holes” sometimes visible inside their cytoplasm (Fig 4e). Flagella (diameter: ~ 0.009 μm) could also be detected all around bacteria (Figs 4b-4d and 4f). Additionally, extruded trichocysts (extrusive organelles typical of *Paramecium*), presumably included in *Paramecium* homogenate and ingested by treated planarians, were easily recognizable inside intestinal cell phagosomes (Fig 4g).

A similar scenario was observed 7 days after feeding, when most of the bacteria occurred enclosed inside planarian enterocyte phagosomes (Figs 5a and 5b); some appeared degraded (not shown), while many others did not show degraded conditions and could survive in the phagosomes (Figs 5a and 5b). In few cases, bacterial-like circular-ovoid shapes were also observed in the cytoplasm of intestinal cells, outside from phagosomes (Fig 5c).

No bacteria were ever observed in tissues other than intestine in treated planarians; similarly, no bacteria were observed in tissues of investigated TEM-processed control animals (S3 Fig).

## Discussion

### Trans-infection experiments

The alphaproteobacterium “*Ca*. Trichorickettsia mobilis subsp. hyperinfectiva”, previously described in the cytoplasm of *P. calkinsi* [63], was retrieved in this study also in the macronucleus of the ciliate *P. multimicronucleatum* strain US_Bl 16I1 s. “*Ca*. Trichorickettsia mobilis” up to now has been exclusively retrieved as an endosymbiont of ciliates belonging to the genera *Paramecium* and *Euplotes* [28, 63]. At present, three subspecies of this RLO endosymbiont have been identified on molecular basis [63]; a comparison among them in the light of the present findings, which suggest a certain morphological plasticity of this bacterium, is presented in Supplementary Material.

The aim of the present paper was to verify the potential trans-infection of the *Rickettsia*-related macronuclear endosymbiont “*Ca*. Trichorickettsia mobilis subsp. hyperinfectiva” of the ciliate *P. multimicronucleatum* strain US_Bl 16I1 to the metazoan model planarian *D. japonica*. There are several studies reporting on the host/symbiont relationships of different *Paramecium* species with different *Rickettsiales*; *P. multimicronucleatum* lies in this ciliate selection, and is a rather a common species, sharing the freshwater habitat with planarians. Thus, it was chosen as donor in trans-infection experimental context as, in our opinion, it can be considered a suitable candidate as putative environmental vector for RLOs. Additionally, the biology of this ciliate host is well-known, and the strain US_Bl 16I1 could be comfortably cultivable under laboratory conditions using flagellates instead of bacteria as food source.

According to the present findings, the trans-infection experiments were successful, i.e. they showed the capability of “*Ca*. Trichorickettsia mobilis” to enter the tissues of planarians. Indeed, in the intestine of planarians, previously antibiotic-treated to avoid bacterial contamination and fed on liver paste enriched with pellet of ciliate homogenate (including “*Ca*. Trichorickettsia” symbionts), we could observe up to day 7 after feeding the presence of flagellated bacteria with a morphology and a size fully resembling those of the RLO endosymbiont of *P. multimicronucleatum* US_Bl 16I1. In our TEM experiments, besides undigested bacteria enclosed in phagosomes, in few cases circular-ovoid shapes, resembling “*Ca*. Trichorickettsia” bacteria, were observed in the cytoplasm of planarian intestine cells, free from phagosomal membrane (Figs 4 and 5).

On the contrary with respect to treated animals, in TEM preparations of controls (i.e. antibiotic-treated planarians fed on plain liver paste) no bacteria were found (S3 Fig).

These results were confirmed and supported by PCR analysis and sequencing of obtained amplicons: the DNA of “*Ca*. Trichorickettsia mobilis” was recovered in treated planarians up to day 7 after feeding while in control animals no RLO DNA was amplified and detected. On the other side, no morphological or behavioural alterations were observed in the planarians

Our findings are in line with those by [99]. These authors studied the phylogenetic identities of digestion-resistant bacteria that could survive starvation and form relatively stable associations with some marine and freshwater ciliate species, and demonstrated that the classes *Alphaproteobacteria* (which includes the order *Rickettsiales*) and *Gammaproteobacteria* are prevalent as digestion-resistant bacteria; from this study a putative significant role of secretion systems in promoting marine protist-bacteria associations resulted as well.

In our experiments, after being ingested by planarians, the bacteria were observed enclosed inside phagosomes of intestinal cells. This also occurred in previous experiments investigating the resistance of planarians to infection by bacterial strains pathogenic for humans and other metazoans [67]. In that research, planarians could eliminate most of the phagocytised bacterial strains within 3-6 days post-feeding thanks to 18 resistance genes, such as *MORN2*, so that the authors suggested that planarians can be considered a model to identify innate resistance mechanisms. Thus, the evidence we obtained that the “*Ca*. Trichorickettsia” endosymbionts of *P. multimicronucleatum* US_Bl 16I1 are still detectable in planarian intestine enterocytes inside and outside phagosomes up to 7 days after feeding could possibly indicate the capability of “*Ca*. Trichorickettsia mobilis subsp. hyperinfectiva” to avoid typical defence mechanisms exploited by planarians.

TEM observations indicated that in some cases *P. multimicronucleatum* RLO could possibly occur outside digestive vacuoles, whose membrane often appeared fractured. Thus, we can hypothesize that this bacterium can transitionally enter and perhaps survive within planarian tissues, although present results do not allow yet to drive unambiguous conclusions on this regard. Interestingly, similarly to some *Gammaproteobacteria* such as *Rheinheimera* sp. strain EpRS3 (*Chromatiaceae*), capable of escaping from phagosomes of the ciliate *Euplotes aediculatus* when fed on the bacterium plus its culture medium [100], *Rickettsiaceae* are already known for their ability to escape the host vacuolar membrane, residing freely in the host cytoplasm, where they may exploit host cytoskeleton for movement [101–103, 12, 17].

Another interesting parallelism might be made with the so-called “eta” particles (i.e. intracellular bacteria) hosted in the cytoplasm of the ciliate *Euplotes crassus* killer cell strain M on *E. crassus* sensitive cell strain 21A7 fed on strain M homogenate [104]. In that study, “eta” particles conferred ciliate strain 21A7 a transitory resistance to killing effect during their permanence as free entities in its cytoplasm: indeed, after being collected into the digestive vacuoles, “eta” particles were pushed towards the periphery of the vacuole and those still not digested were capable of escaping into ciliate cytoplasm through vacuole membrane rupture.

Unfortunately, just as in the studies by Chiellini et al. [100] and Verni et al. [104], we are currently not able to provide here clues on the potential bacterial mechanisms involved in planarian phagosome escaping.

To the best of our knowledge, this is the first time that a set of experimental bioassays was performed to verify the transmission of a “true” and ascertained *Rickettsiales* bacterium from an infected protist to an uninfected metazoan of the same aquatic environment (freshwater). On the other side, in the past, efforts have been put to experimentally verify the transmission of morphologically RLOs among aquatic organisms. For example, Nunan et al. [105] performed bioassays to verify the transmission of the infection between two species of commercially farmed shrimps, i.e. the infected *Penaeus monodon* and the specific pathogen-free *Penaeus vannamei*, with the aim of investigating the suspected causative agent of severe mortality in farms where those organisms are in co-culture (grow-out ponds). The bioassays were performed both via injection of infected shrimp homogenate into uninfected shrimps and per *os* exposure of uninfected shrimps to tissue taken from infected shrimps. Only injection bioassays were successful leading to an infection, while per *os* infection failed. Among different possible reasons for this negative result those authors cited the potential need for a vector to disseminate the disease. According to our findings, ciliates could represent suitable vectors in this kind of situation. In our experiments, as distinct from Nunan et al. [105], we chose only to perform the bioassay per *os* exposure of animals instead of injection into the planarian intestine. Endosymbionts, separated from ciliate cells through cell rupture and centrifugation, could be easily mixed with planarian food and seeded on the bottom of a Petri dishes so as to allow animals to reach it and feed. As we dealt with endosymbionts, which are present in limited numbers inside their ciliate host, we chose to treat the planarians with cell mass culture homogenate instead of adding living ciliates to planarians food. This allowed processing of as many ciliates as possible to maximize the probability of endosymbiont ingestion by the animals, thereby increasing the chance of detection of successful trans-infection via PCR and TEM-based approaches.

We believe that our findings may offer intriguing insights when considered from several points of view, such as concerning the pathologies caused by *Rickettsiales* or RLOs occurring in fish farms or in the wild, which might have ciliates or other protists as putative vectors. Although there is still a need for further investigations on this topic to expand its implications, we think that our study can serve as basis for conceiving long-lasting experiments aiming to better understand whether “*Ca*. Trichorickettsia mobilis”, as well as other *Rickettsiales* symbionts of protists, can be able to survive longer and potentially replicate in tissues of planarians and other aquatic Metazoa, and whether these RLOs may have some impact on the recipient host health.

## Compliance with Ethical Standards

### Ethics Approval

This study does not contain any studies with human participants performed by any of the authors. All procedures performed in studies involving animals were in accordance with the ethical standards of the institution or practice at which the studies were conducted.

### Conflict of Interest

The authors declare that they have no conflict of interest.

## Acknowledgments

The authors wish to thank Francesco Paolo Frontini, Fabrizio Erra, and Federica Vantaggio for ciliate culturing and *in vivo* processing; Claudio Ghezzani and Simone Gabrielli for assistance with TEM material; Simone Gabrielli for help with graphic artwork, Thomas Berendonk for the opportunity to collaborate on *Paramecium*. Financial support: project PRA_ 2016_58, University of Pisa to GP; RFBR grant N° 18-04-00562-a to ES. All authors critically read and approved the manuscript.

## Author Contributions

Conceptualization: GP, FV, AS, LM, LR.

Formal analysis: LM, AS, LR, FS, MC.

Funding acquisition: GP, ES.

Investigation: LM, AS, LR, FS.

Project administration: GP.

Supervision: GP, FV.

Visualization: LM, AS.

Writing – original draft: LM, AS, LR, MC.

Writing – review & editing: FV, GP, FS, ES, GDG, SIF, SK.

## S1_Supporting information

**S1 Fig. Light microscope observation of *P. multimicronucleatum* strain US_Bl 16I1**. (**a-c**) *In vivo* specimens; (**d**) Feulgen stained cell. Ma, macronucleus. (**c**) the cytostome (C) at a higher magnification with trichocysts (T) inserted in the cortex. (**d**) Feulgen staining highlights the Ma and the three micronuclei (mi). Scale bars stand for 10 μm.

**S2 Fig. TEM pictures of *P. multimicronucleatum* strain US_Bl 16I1**. (**a, b**) cortex with trichocysts inserted (T) in resting state (**a**) and about to extrude (**b**); m, mitochondria. (**c, d**) macronucleus (Ma) encircled by a layer of rarefied material (asterisk) with endosymbionts (arrows) and nucleoli (n); Ph, phagosomes. (**e**) Endosymbionts inside the Ma, a transversally sectioned T occurs near Ma membrane. (**f**) Ma portion and one of the three micronuclei (mi). Scale bars stand for 1 μm.

**S3 Fig. TEM pictures of the intestine of *D. japonica* specimens fixed after feeding with plain liver paste (control animals)**. (**a**) day 1 after feeding; (**b**) day 7 after feeding. L, lumen of planarian intestine. Scale bars stand for 1μm.

